# Altered theta distribution and coherence during set-shifting in older age

**DOI:** 10.64898/2026.01.27.701912

**Authors:** Margarita Darna, Anni Richter, Jens-Max Hopf, Constanze I. Seidenbecher, Björn H. Schott

**Affiliations:** Leibniz Institute for Neurobiology (LIN), Magdeburg, Germany; Department of Medical Psychology, Medical Faculty, Otto-von-Guericke University Magdeburg, Germany; German Center for Mental Health (DZPG), partner site Halle-Jena-Magdeburg; Center for Intervention and Research on adaptive and maladaptive brain Circuits underlying mental health (C-I-R-C), Halle-Jena-Magdeburg; Center for Behavioral Brain Sciences (CBBS), Magdeburg, Germany; Department of Psychiatry and Psychotherapy, Medical Faculty, Otto-von-Guericke University Magdeburg, Germany; German Center for Neurodegenerative Diseases (DZNE), Göttingen, Germany; Department of Psychiatry and Psychotherapy, University Medical Center Göttingen, Göttingen, Germany; University Clinic for Neurology, Otto-von-Guericke University Magdeburg, Germany

**Keywords:** Aging, set-shifting, EEG, cognitive flexibility, theta oscillations, coherence

## Abstract

Cognitive flexibility is an executive function that enables adapting behaviour to a changing environment and is thus critical for daily life. The degree of its preservation upon healthy aging and the neural mechanisms underlying it are still a matter of debate. To investigate the electrophysiological correlates of cognitive flexibility in older age, we measured cognitive flexibility in 99 young (24.75 ± 4.45 years) and 83 older adults (69.19 ± 6.25) using electroencephalography (EEG). Compared to young adults, older adults showed a more conservative response pattern with longer reaction times, but lower error rates (speed-accuracy tradeoff). In the EEG, both age groups exhibited increased theta-power during set-shifting, with a fronto-central peak in the young, but a more fronto-lateral topography in older adults. Importantly, both groups displayed increases in theta coherence and global efficiency during set-shifting, but coherence modulations were restricted in frontocentral areas in the young but were diminished and distributed across the scalp in the older. Better set-shifting performance was most strongly associated with high coherence and global efficiency irrespective of age group. These results point to an age-related change of cortical processing underlying cognitive flexibility which involves the employment of more distributed neural resources for successful task completion.

## 1. Introduction

Cognitive flexibility is an executive function that enables individuals to adapt their behaviour to a changing environment (Diamond, 2013). Its importance becomes evident in its associations with salutogenetic factors, such as trait resilience (Genet and Siemer, 2011), mindfulness (Moore and Malinowski, 2009) and social adjustment (Chen et al., 2024). In older age, in particular, cognitive flexibility has been linked to overall quality of life (Davis et al., 2010) and to planning and fluid intelligence in the presence of mild cognitive impairment (MCI) (Corbo et al., 2024). Despite its importance, age-related changes in cognitive flexibility at both the behavioural and the neural level are thus far insufficiently understood. The present study was conducted with the goal to evaluate age-related changes of set-shifting performance and their association with electrophysiological measures of brain activity.

In laboratory settings, cognitive flexibility is often operationalized as task-switching and set-shifting performance using paradigms, which require participants to shift their attention between different tasks, rules or dimensions. Performance is then assessed by comparing trials with and without shifts. Shift trials can include intra-dimensional (ID) or extra-dimensional (ED) shifts (Watson et al., 2006). In ID shifts, the new correct feature belongs to the previously correct dimension, whereas in ED shifts, the correct feature belongs to a different dimension. Deficits in cognitive flexibility usually appear as switch costs, most notably increased reaction times and/or error rates compared to trials without shifts (Wylie and Allport, 2000), and with perseverative errors, that is, when a previously correct response pattern is kept on being employed after the shift required a change (Heaton et al., 1993).

Studies on behavioural manifestations of age-related changes in cognitive flexibility have thus far yielded inconsistent results. While several studies report deficits reflected by increased switch costs (Cepeda et al., 2001; Kray et al., 2002; Meiran et al., 2001) and more perseverative errors (Haaland et al., 1987; Zelazo et al., 2004), others describe an unaltered (Falkenstein et al., 2001; Karayanidis et al., 2011; Kolev et al., 2005; Kray and Lindenberger, 2000) or even improved performance compared to younger adults (Kray, 2006). Only few studies in older age have assessed the impact of different set-shifting demands, most notably ID versus ED shifts, with the latter engaging more neural resources (Watson et al., 2006). Again, some studies reported older adults to show increased difficulties during ED shifts (De Luca et al., 2003; Owen et al., 1991; Zelazo et al., 2004) but in our previous study (Darna et al., 2025), we found no evidence of higher ED switch costs in older adults.

At the neural level, studies on error processing (Kolev et al., 2024) and motor coordination (Yordanova et al., 2020) point to a functional reorganization of frontal networks in older age. In the context of cognitive flexibility, evidence for such reorganization has been found, in form of an age-related topographical shift of theta (4-8 Hz) amplitude modulations from parietal to fronto-temporal sensors (Huizeling et al., 2021). Other studies have reported attenuated frontal theta amplitude modulations in older adults during reversal learning (Küçük et al., 2023) and during both ID and ED shifts (Darna et al., 2025).

At the network level, lower theta coherence across brain regions has been observed in older adults during task-switching (Dias et al., 2015). Finally, increased global efficiency – a network-based measure reflecting the efficiency of information transmission across a network (Latora and Marchiori, 2001) – has been linked to higher cognitive demands (e.g. Cohen and D’Esposito, 2016) and is hypothesized to act as a mediator for executive functions in older age (Li et al., 2020), despite its age-related decrease (Ajilore et al., 2014; Li et al., 2020).

Taken together, these theta-band neural signatures refer to potential mechanisms underlying set-shifting performance in older age. To investigate their role, we applied the ID/ED set-shifting task (IDED) (Darna et al., 2025; Oh et al., 2014) and recorded electroencephalographic brain activity (EEG). We chose the IDED, as it enabled us to isolate the different types of shifts (i.e., ID and ED), an aspect that is usually absent in other studies of cognitive flexibility.

We hypothesized that theta measures would be differently modulated in older individuals compared to young. More specifically we hypothesized that older participants would exhibit a) lower theta power and reduced theta power modulation in both ID and ED set-shifts, b) overall lower theta coherence between channel pairs c) lower global efficiency and reduced modulation of global efficiency. We also investigated the relationship between performance measures and theta measures using a linear mixed effect model approach and particularly explore age related changes.

## 2. Materials and Methods

### 2.1 Participants

Recruited participants were between the age of 18 to 35 to be included in the young age group and at least 60 years old to be recruited in the older age group. The Mini-International Neuropsychiatric Interview (M.I.N.I; Sheehan et al., (1998); German Version by Ackenheil et al., (1999)) and a standardized health questionnaire (Richter et al., 2011) were conducted to exclude past or present manifest neurological or psychiatric disorders, substance abuse, use of neurological or psychiatric mediation or serious medical conditions (e.g., heart failure NYHA stage III or IV, metastatic cancer, or diabetes mellitus with complications). Additionally, only right-handed individuals with fluent German language skills participated. The study was approved by the Ethics Committee of the Faculty of Medicine at the Otto von Guericke University of Magdeburg. All individuals gave written informed consent in accordance with the Declaration of Helsinki (World Medical Association Declaration of Helsinki, 2013) and participation was compensated financially or with course credit, depending on participants’ preference.

Additional questionnaires to evaluate cognitive status were the German version of the crystallized intelligence test “Mehrfachwahl-Wortschatz-Intelligenztest B” (MWT-B; Lehrl, 1999; Lehrl et al., 1995). Older participants also completed the Mini Mental State Examination (MMSE; (Folstein et al., 1975)).

In total, 188 participants completed the study, out of which three older adults were excluded due to the self-reported inability to understand the task and three additional participants (2 young, 1 older) were excluded as they were identified as extreme outliers in the IDED task, as indicated by mean reaction time or error rates (criterion: 3^rd^ quantile + 3*interquartile range)(see Table S2). The final sample comprised 182 participants, including 99 young (age: 24.76 ± 4.44 years) and 83 older (age: 67.53 ± 12.36 years). Participant groups did not differ with respect to sex distribution and education years and old participants displayed a significantly higher MWT-B score as seen in our previous study (Darna et al., 2025).

A subset of the participants took part in an extended neuropsychological testing battery to investigate further psychological constructs such as attention, short- and long-term memory, working memory and executive functions (as described in (Richter et al., 2023); for results see Table S1 in supplementary material). One task of interest for this study is the flexibility subtest of the Test Battery for Attention (TAP) (Zimmermann and Fimm, 1992). All participants also completed the attentional set-shifting task (ASST)(Sahakian and Owen, 1992) as described in Darna et al., (2025). The results of these two set-shifting tasks are presented in Table 2 and supplementary Figure 1.

**Table 1.** Participant summary statistics.

|  | **Young**<br>**M ± SD [N]** | **Older**<br>**M ± SD [N]** | **Statistics** |
| --- | --- | --- | --- |
| N | 99 | 83 |  |
| Age | 24.76 ± 4.44 [99] | 67.53 ± 12.36 [83] | ***W* = 8217, *p* < .001** |
| Sex | 37 M, 64 F | 39 M, 44 F | $X^2_1 = 2.07$, *p* = .150 |
| Education (years) | 15.82 ± 2.81 [97] | 15.30 ± 2.23 [81] | *W* = 3576, *p* = .301 |
| MWT-B | 26.01 ± 3.69 [98] | 31.68 ± 2.69 [82] | ***W* = 7165, *p* < .001** |
| MMSE | - | 28.51 ± 1.12 [83] | - |

M: mean; SD: standard deviation; N: sample size; F: female; M: male; *t*: Welch two-sample *t*-test; $X^2$ : Pearson’s Chi-squared test; *W*: Wilcoxon rank sum test; MWT-B: Mehrfachwahl-Wortschatz-Intelligenztest B; MMSE: Mini Mental State Examination.

**Table 2.** Behavioural flexibility tasks – summary results in young and old.

|  | **Young**<br>**M ± SD** | **Older**<br>**M ± SD** | **Statistics** |
| --- | --- | --- | --- |
| **Flexibility subtest from TAP** |  |  |  |
| N | 53 | 63 |  |
| Error rate (%) | 3.97 ± 4.13 | 11.06 ± 13.78 | ***W* = 2220.5, *p* = .002** |
| RT (ms) | 1071.42 ± 199.98 | 1918.74 ± 590.94 | ***W* = 3208, *p* < .001** |
| **ASST** |  |  |  |
| N | 100 | 80 |  |
| Overall survival probability<br>[N-failed] | 80 % [20] | 43.80 % [45] | ***X*<sup>2</sup> = 26.40, *p* < .001** |
| *First Reversal Stage (CDR)* |  |  |  |
| N | 99 | 77 |  |
| RT (ms) | 886.73 ± 395.32 | 1108.83 ± 406.49 | ***W* = 5327, *p* < .001** |
| Trials-to-criterion | 10.91 ± 3.55 | 13.62 ± 8.89 | ***W* = 4542, *p* = .023** |
| *First ID Stage (IDS1)* |  |  |  |
| N | 99 | 76 |  |
| RT (ms) | 895.24 ± 390.02 | 1152.50 ± 355.68 | ***W* = 5617, *p* < .001** |
| Trials-to-criterion | 9.23 ± 2.27 | 10.55 ± 5.47 | *W* = 4279, *p* = .095 |
| *ED stage (ED)* |  |  |  |
| N | 80 | 45 |  |
| RT (ms) | 1033.84 ± 327.83 | 1391.79 ± 385.72 | ***W* = 2146, *p* < .001** |
| Trials-to-criterion | 17.55 ± 8.14 | 25.29 ± 11.21 | ***W* = 2020, *p* < .001** |

M: mean; SD: standard deviation; N: sample size; *X*<sup>2</sup>: Kaplan–Meier survival comparison using the log-rank test; *W*: Wilcoxon rank sum test; CDR: Compound Discrimination Reversal; IDS1: Intra-dimansional shift 1

**Figure 1:**
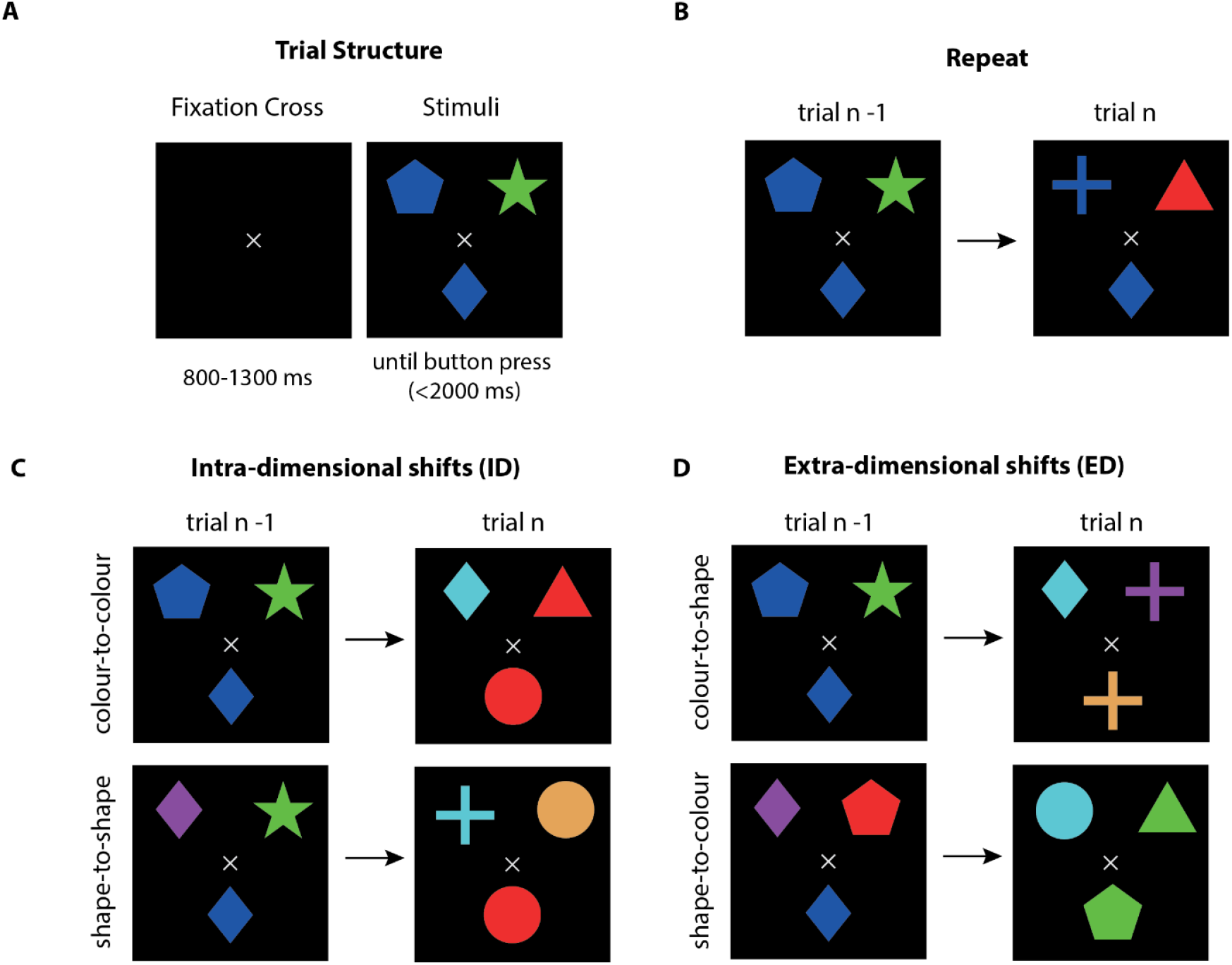
The IDED task (adapted from Darna et al., 2025). **A:** Trial structure. After the presentation of the fixation cross for a pseudorandomized interval between 800 ms to 1300 ms, the stimuli (top) and target (bottom) appear. These are visible until a button press occurs or until 2000 ms have elapsed; **B:** Example of a repeat trial. Here, the target (here blue diamond) and the matching rule (match blue colour) remain the same from trial to trial; **C:** Example of intra-dimensional shifts (ID). Here, the matching rule changes from colour to colour (top row, e.g. blue-to-red) or from shape to shape (bottom row, e.g. diamond-to-circle). **D:** Example of extra-dimensional shifts (ED). Here, the matching rule changes from colour to shape (top row, e.g. blue-to-cross) or from shape to colour (bottom row, e.g. diamond-to-green).

### 2.2 Cognitive Flexibility Task

All participants completed the IDED, which has previously been described in detail (Darna et al., 2025; Oh et al., 2014) and is only briefly presented here (including some minor modifications from the previous version)(Figure 1). The paradigm was executed using the Psychtoolbox (Brainard and Vision, 1997; Pelli, 1997) running on MATLAB R2021b (The MathWorks Inc., 2021, Natick, MA). Each trial began with a fixation cross presented for 800 to 1300 ms (jitter generated with uniformly distributed pseudorandom numbers in MATLAB), followed by the presentation of the target below the fixation cross and the stimuli above. Participants were asked to indicate the stimulus matching the target via a mouse click (left or right). The matching features could be the colour or the shape of the target. After the button press or a maximum response time of 2000 ms, the stimuli disappeared and a new trial began. Participants received no feedback on the correctness of their response.

The task consisted of 100 ID and 100 ED trials each, in which the matching criterion was changed within a dimension or across dimensions respectively. After each shift trial, randomly 2 to 7 trials with no set-shift followed, in which the target and matching criterion remained the same (hence termed repeat trials). We only evaluated the second repeat trials after each set-shift resulting in 200 repeat trials (as done in: Darna et al., 2025).

Participants were explicitly instructed to respond as fast as possible, and a practice run of 20 trials was performed before the experiment, during which participants received feedback on the correctness of their responses. Here, we also ensured that the participants could differentiate the colours of the stimuli.

The session consisted of 6 blocks with self-paced breaks in between and lasted approximately 50 to 60 minutes. The practice trials and repeat trials before the first shift after a break were excluded from analysis. The recording took place in a dimmed, electrically shielded room. Participants were sitting at a distance of 110 cm away from the 32” presentation monitor with a resolution of 1920×1280 pixels and a refresh rate of 120 Hz. Mouse button responses were performed with the right hand.

### 2.3 Statistical Analysis of behavioural measures

We evaluated performance in the IDED in RStudio R 4.5.1 (R Core Team, 2022; R Studio Team, 2020) using three distinct measures:

1. Reaction time (RT) was defined as the time taken to correctly respond after stimulus presentation. For RT analysis we excluded short RTs due to anticipatory responses (< 150 ms) and evaluated the median RT for each condition as RTs displayed a right-skewed distribution (mean skewness for all trials: 0.91 ± 0.26).
2. Interquartile range of RTs (IQR) was evaluated to represent intertrial variability in performance.
3. Error rates were computed as the percentage of trials with incorrect, early (<150 ms) or no responses in each condition.

To evaluate switch costs in set-shifting performance we calculated them as the difference between ID and repeat and ED and repeat for RTs, IQRs and error rates as follows:

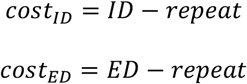

This resulted in two switch cost estimates per behavioural measure used as dependent variables for six separate robust analyses of variance (ANOVA) with the package WRS2 (Mair and Wilcox, 2014) with the factors age group (young vs. older; between group) and condition (ID vs ED; within group). Whenever an interaction effect was significant post-hoc *t*-tests on the trimmed means were conducted and a robust cohen’s *d* on 20% trimmed means was calculated as effect size.

To evaluate global switch costs, we calculated a composite performance z-score as the sum of the standardized switch scores for each performance measure according to:

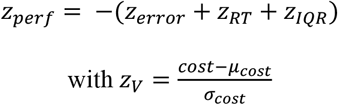

where *cost* represents the individual switch cost of an individual performance measure (Darna et al., 2025; Dias et al., 2015; Liesefeld and Janczyk, 2019) and *μ* and *σ* represent the average and standard deviation, respectively, collapsed across conditions and age groups. Here, high z-scores indicate lower global switch costs, in other words, a better performance in the respective condition. Importantly, *μ* and *σ* were balanced to account for the different sample sizes of the two age groups. Here, we again performed a robust ANOVA and post-hoc *t*-tests as described above to evaluate the effect of age group and condition on the z-score.

### 2.4 EEG Acquisition and Pre-processing

We recorded the EEG at 1000 Hz sampling rate using 64 active electrodes (Brain Products GmbH, Gilching, Germany) of the actiCAP layout (EASYCAP) and an additional ground electrode at AFz. Online reference was the FCz. Four EOG electrodes were recorded for detection of (micro)saccades and blinks. These were: a horizontal EOG of the left eye (HEOGL), horizontal EOG of the right eye (HEOGR), superior vertical EOG of the left eye (VEOGS) and inferior vertical EOG of the left eye (VEOGI). The EEG was recorded with BrainRecorder from Brain Products and port event signalling was conducted the Mex-File Plug-in IO74 (https://apps.usd.edu/coglab/psyc770/IO64.html). Electrode impedances were kept below 10kΩ with electrolyte gel.

The first pre-processing steps were conducted using Brain Vision Analyzer 2.2 (Brain Products GmbH, Gilching, Germany). First, data were re-referenced to the average of all EEG channels and the old online reference was reused as a channel (FCz). Next, new EOG channels were calculated as followed: horizontal EOG: HEOG = HEOGR – HEOGL, vertical EOG:VEOG = VEOGI – VEOGS and radial EOG: REOG = ((HEOGR + HEOGL + VEOGI + VEOGS)/4) – Pz. The data were then filtered with a zero-phase shift Butterworth low-pass filter at 250 Hz with the order of 2. Artifacts were marked using a semi-automatic raw data inspection with a maximum allowed voltage step of 70 μv/ms (Gradient; marked 400 ms before and after the event) and lowest allowed activity in intervals of 0.5 μV for 100 ms (Low Activity; marked 200 ms before and after the event). Here, channels with more than 20% marked datapoints were visually inspected by a trained observer (author MD) for future channel interpolation. Next, eye blinks were detected via automatic raw data inspection on the VEOG channel. Here, all intervals were marked where the allowed difference of values in an interval length of 200 ms exceeded a value 0f 100 μV (Max-Min; marked 100 ms before and after the event). The continuous data sets were then exported for further processing in MATLAB.

Within the MATLAB environment (R2024b), we used EEGLAB to import the data (Delorme and Makeig, 2004; Version 2024.0). Bad segments that were marked before, were now removed from the data and blinks were added as events in the structure. The EOG channels of individual subjects (n = 8) with line noise in EOG were filtered with a Butterworth 2nd order notch filter from 48 to 52 Hz. Then, we extracted stimulus-locked trial epochs from −1500 ms to 3000 ms for the trials of interest (repeat, ID and ED) that were followed by a correct response. Here, incorrect responses also included trials without responses and trials with premature responses (< 150 ms), representing anticipatory responses. Participants with fewer than 30% trials remaining within each condition were excluded from further analysis (n young = 7; n older = 24), resulting in 92 young and 59 older participants included in the analysis. The epochs were baseline corrected with the average obtained from the time window between −250 to 0 ms. Saccades, microsaccades and muscle artifacts were detected and corrected with the EEGLAB plug-in microDetect (Craddock et al., 2016) and an Independent Component Analysis (ICA; FASTICA; http://research.ics.aalto.fi/ica/fastica/)(for details see supplementary material)

After ICA, individual noisy channels of the previously inspected datasets (10 subjects; see table S3) were interpolated in EEGLAB with spherical spline interpolation (*λ =* 0, m = 4, n = 7).

The datasets obtained from this step, were processed separately in each one of the following three sections.

### 2.5 Theta Power Analysis

For time-frequency analysis, data were first converted to Fieldtrip format using the “eeglab2fieldtrip” function in MATLAB. In Fieldtrip (Oostenveld et al., 2011; Version 20220104), frequency power from 1 to 100 Hz was calculated using Morlet wavelets with a width of 7 cycles and length equal to 3 standard deviations of the implicit Gaussian kernel, using a sliding time window in steps of 10 ms. Oscillation power was then baseline-corrected and normalized to DB (baseline range: −300 to −100 ms before stimulus presentation) as follows:

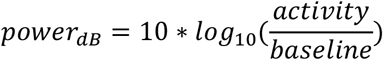

We then extracted the mean normalized power of the theta frequency band (4-8Hz) across all channels in the time window between −300 ms and 1000 ms for each participant and condition and conducted a factor analysis (FA) in the temporal domain on the individual age groups. FA was performed as described by Scharf et al. (2022). Briefly, the Empirical Kaiser Criterion was employed to determine the number of factors to retain (eigenvalue > 1) and the unrotated factor loadings of the exploratory factor analysis were first estimated using custom R code and (Braeken and Van Assen, 2017). Next, the Geomin rotation (Yates, 1987) was applied, an oblique rotation technique that minimizes the complexity of the loading matrix at the cost of allowing some degree of correlation between factors. The resulting components were then evaluated based on their maximum peak latencies and overall topography. Two components were chosen for further analysis, as they explained the highest variance, displayed the highest unstandardized loadings and their peak was around the post-stimulus time-range of interest based on our previous study (Darna et al., 2025): an early theta component with a peak latency at 430 ms in the young group and 440 ms in the older age group and a late theta component with a peak latency at 680 ms in the young and 700 ms in the older group. To estimate factor scores, we extracted factor-wise reconstructed theta power time series across all electrodes. The peak-latency amplitudes of the respective reconstructed data were used in the statistical analyses.

### 2.6 Coherence and Global Efficiency

To circumvent the common reference problem (Bastos and Schoffelen, 2016), we computed Current Source Density estimations (CSD) from the raw EEG data using the CSD Toolbox (Kayser and Tenke, 2006; https://psychophysiology.cpmc.columbia.edu/software/csdtoolbox/). Coherence was calculated using a bootstrap procedure with 100 runs to account for the sample size bias resulting from different numbers of trials across conditions (Bastos and Schoffelen, 2016). In each bootstrap run, we randomly sampled 30 trials from each condition and calculated coherence values as follows. Spectral decomposition was performed using Morlet wavelets as described above, with the frequencies of interest now ranging from 1 to 10 Hz. We used the resulting Fourier spectra in the subsequent connectivity estimation with Fieldtrip’s,*ft_connectivityanalysis’*. Here, we computed the imaginary coherence between all pairs of channels over the time window from −300 ms to 1000 ms. The raw coherence values across all electrode pairs (*n*_*pair*_ = 2080), was input in a temporal PCA to identify components of interest as described in the section above. The Geomin rotation of the obtained principal components with all conditions did not converge, we thus calculated the temporal PCA with the set-shifting contrasts ID minus repeat and ED minus repeat. Here, we again identified two components of interest as they displayed a peak latency around the same time windows as the components obtained from theta power. The peak-latency amplitudes of the respective reconstructed components were used in the statistical analyses.

Additionally, we extracted the global efficiency for each participant and condition as a network-based measure reflecting the efficiency of information transmission across the entire network (Latora and Marchiori, 2001) using the Brain Connectivity Toolbox (Rubinov and Sporns, 2010). Global efficiency was estimated within two time-windows reflecting the time range of the components obtained from the previous sections:

- early global efficiency was estimated as the mean within the time window of 410 to 510 ms, and
- late global efficiency was measured between 615-715 ms.

### 2.7 Linear Mixed Effects Model

To evaluate whether the theta measures could predict behaviour in the IDED we first standardized the theta measures by subtracting the mean and dividing by the standard deviation across all participants. We then fitted a linear mixed-effects model on the data with z-score as dependent variables and the amplitude of the early theta component (theta_Early), late theta component (theta_Late), early coherence (coh_Early), early global efficiency (gl_Early), late global efficiency (gl_Late) as independent variables. Late coherence was not included as a parameter as it displayed no significant modulation during set-shifting (see results 3.3). The model also included the interaction of these variables with age group and condition and a random intercept for each participant to account for repeated measures as such:

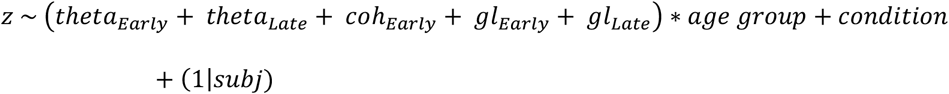

where ‘subj’ refers to the individual participant pseudonym.

To identify the best fitting model, we performed an all-subsets model selection using the ‘dredge()’ function from the MuMIn package in R and ranked models by their Akaike Information Criterion (AIC). The three best-fitting models were extracted and are being described in the results.

### 2.8 Statistics

For all statistical tests reported, the alpha significance level was set to *p* = 0.05. ANOVA results are reported with the obtained *F* value, *p* value and partial eta squared (*η*_*p*_^2^) as measure of effect size. T-tests are reported with the obtained *t* value, *p* value adjusted for multiple comparisons with the Bonferroni-Holm correction, and Cohen’s d as effect size. Statistical significance is reported in the graphs as followed: * for *p*≤0.05, ** for *p*≤0.005 and *** for *p*≤0.001. Average values in text are reported with standard deviations in the form: mean ± standard deviation, unless otherwise specified.

## 3. Results

### 3.1 Behavioural Results of the IDED Task

Overall, the average median RT was 533 ms ± 133 ms, with older participants reacting more slowly than the young (young: 469 ms ± 42 ms; older: 706 ms ± 109 ms; *t*_102_ = −18.77, *p* < 0.001, *d* = −2.88). A robust ANOVA of the switch costs (difference in RT between ID or ED from repeat) revealed a significant interaction between age group and condition (*F*_1,78.05_ = 18.26, *p* < 0.001). Both age groups showed increased RT switch costs in ED compared to ID trials (young: *t*_*60*_ = −18.23, *p* < 0.001, *d* = −0.89; older: *t*_*50*_ = −11.74, *p* < 0.001, *d* = −0.9), but older participants showed significantly higher RT switch costs than the young in the ED condition (ID: *t*_*76.63*_ = −1.89, *p* = 0.062, *d* = −0.23; ED: *t*_64.03_ = −4.18, *p* < 0.001, *d* = −0.67)(**Figure 2**A). A main effect of condition was also found (*F*_1,78.05_ = 309.43, *p* < 0.001), where switch costs were significantly higher in the ED condition compared to the ID (*t*_*109*_ = −25.44, *p* < 0.001, *d* = −0.85). Finally, there was also a significant main effect of age group (*F*_1,79.13_ = 14.18, *p* < 0.001), with older adults displaying higher RT switch costs compared to young (*t*_*158.76*_ = 3.58, *p* < 0.001, *d* = −0.31).

**Figure 2:**
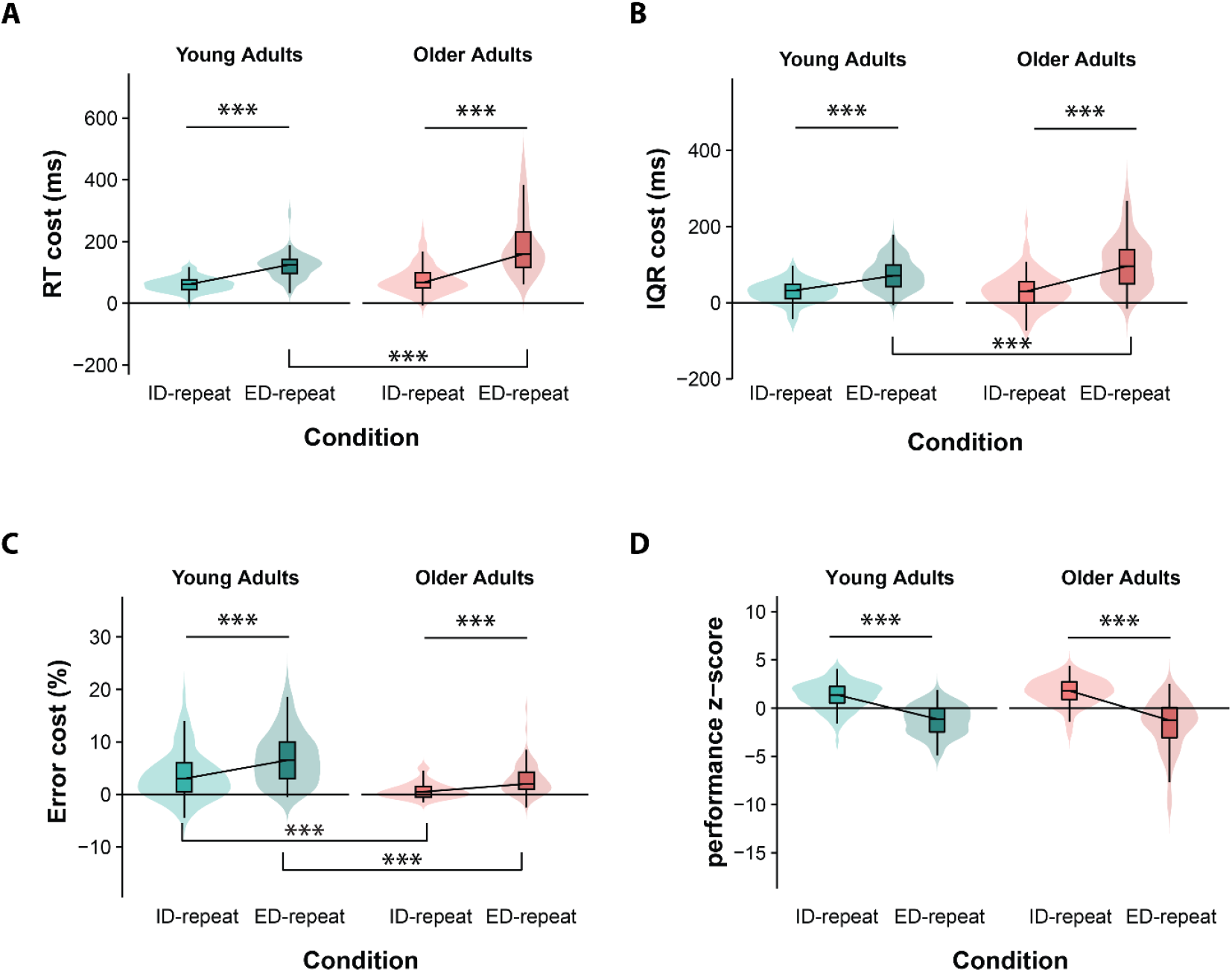
Behaviour in set-shifting. **A:** Mean reaction time (RT) costs as difference between the respective set-shifting condition and repeat. Both age groups displayed increased RT costs in the ED condition compared to the ID condition (young: *t*_*60*_ = −18.23, *p* < 0.001, *d* = −0.89; older: *t*_*50*_ = −11.74, *p* < 0.001, *d* = −0.9). Older adults had higher RT costs in the ED condition compared to young ED: *t*_64.03_ = −4.18, *p* < 0.001, *d* = −0.67); **B:** Mean cost of interquartile range of reaction times (IQR) compared to the repeat condition. Both age groups had increased costs in ED compared to ID (young: *t*_60_ = −9.06, *p* < 0.001, *d* = −0.75; older: *t*_50_ = −8.61, *p* < 0.001, *d* = −0.68). Older adults had higher ED costs compared to young (*t*_81.67_ = −2.76, *p* = 0.007, *d* = −0.31); **C:** Mean error cost compared to the repeat condition. Both age groups again displayed increased error costs in ED compared to ID (young: *t*_60_ = −7.72, *p* < 0.001, *d* = −0.48; older: *t*_50_ = −6.00, *p* < 0.001, *d* = −0.63). Here older adults, however, had lower error costs in both contrasts compared to young adults (ID: *t*_74.15_ = 5.50, *p* < 0.001, *d* = 0.59; ED: *t*_98.03_ = 6.44, *p* < 0.001, *d* = 0.63); **D:** Global performance as indicated by composite z-scores of all behavioural measures. Both age groups display a decrease in z-score in the ED contrast compared to the ID contrast (young: *t*_60_ = −14.86, *p* < 0.001, *d* = 0.85; older: *t*_50_ = −13.27, *p* < 0.001, *d* = −0.94). Importantly, no significant z-score difference was found between the two age groups (*F*_1,111_ = 0.49, *p* = 0.486).

Overall, participants showed an IQR of RTs of 197 ms ± 61 ms, with younger participants exhibiting lower mean IQRs compared to older participants (young: 156 ms ± 29 ms; older: 245 ms ± 54 ms; *t*_121_ = −13.54, *p* < 0.001, *d* = −2.06). A significant interaction between condition and age group was present again (*F*_1,98.03_ = 7.51, *p* = 0.007). IQR costs in the ID condition did not differentiate between young and older participants (*t*_82.41_ = −0.16, *p* = 0.973, *d* = 0.02) but older participants exhibited higher IQR costs in the ED condition compared to the young (*t*_81.67_ = −2.76, *p* = 0.007, *d* = −0.31). Both age groups showed an increase of IQR switch costs from the ID to the ED condition (young: *t*_60_ = −9.06, *p* < 0.001, *d* = −0.75; older: *t*_50_ = −8.61, *p* < 0.001, *d* = −0.68) (**Figure 2**B). The robust ANOVA here again revealed a significant main effect of condition (*F*_1,79.13_ = 14.18, *p* < 0.001), where the ED condition displayed higher switch costs compared to the ID (*t*_109_ = −11.61, *p* < 0.001, *d* = −0.76). Additionally, a significant main effect of age group was found (*F*_1,95.77_ = 4.54, *p* = 0.036). Here, older adults displayed a trend for marginally higher IQR switch costs compared to the younger adults (*t*_150.43_ = 1.83, *p* = 0.069, *d* = −0.17).

On average, participants had very low error rates of 4.54% ± 3.65%, with error rates being lower in older compared to young participants (young: 6.28% ± 3.88%; older: 2.46% ± 1.84%; *t*_145_ = 8.70, *p* < 0.001, *d* = 1.26). The robust ANOVA revealed a significant interaction (*F*_1,105.16_ = 9.30, *p* 0.003), with older participants displaying lower error switch costs in both set-shifting conditions compared to the young (ID: *t*_74.15_ = 5.50, *p* < 0.001, *d* = 0.59; ED: *t*_98.03_ = 6.44, *p* < 0.001, *d* = 0.63). Additionally, young participants showed a smaller effect size regarding switch cost differences between ID and ED (*t*_60_ = −7.72, *p* < 0.001, *d* = −0.48) compared to older participants (*t*_50_ = −6.00, *p* < 0.001, *d* = −0.63) (**Figure 2**C). A significant main effect of condition was also revealed (*F*_1,105.16_ = 95.07, *p* < 0.001), where significant higher switch costs were found in the ED condition compared to the ID (*t*_*109*_ = −8.67, *p* < 0.001, *d* = −0.50).Additionally, we found a significant main effect of age group (*F*_1,87.42_ = 48.18, *p* < 0.001), with older adults displaying lower error switch costs compared to young (*t*_*161.41*_ = 3.35, *p* < 0.001, *d* = 0.61).

Finally, we created a composite z-score reflecting global switch costs in the IDED. A robust ANOVA revealed no significant interaction effect between condition and age group (*F*_1,111_ = 3.08, *p* = 0.082) but a significant within effect (*F*_1,111_ = 380.76, *p* < 0.001). Here, both age groups showed a decrease in z-values from the ID to the ED condition (young: *t*_60_ = 14.85, *p* < 0.001, *d* = 0.85; older: *t*_50_ = 13.27, *p* < 0.001, *d* = 0.94), showing increased global switch costs (worse global performance) in the ED condition compared to ID. No significant between group effect was found (*F*_1,110.93_ = 0.49, *p* = 0.486) (**Figure 2**D).

### 3.2 Theta Band Power

Separate temporal FAs on the two age groups revealed two components of interest: early and late theta. The early theta component in the young group peaked over FCz around 420 ms post stimulus and explained 35.84% of the variance, whereas that of the older group reached a maximum over the F8 channel around 420 ms and explained 25.35% of the variance (**Figure 3**). A mixed ANOVA with the factors condition and age group revealed a significant two-way interaction (*F*_1.91,347.65_ = 12.35, *p* < 0.001, η_p_^2^ = 0.06). Here, the young age group displayed increased component power between all conditions (ID-repeat, *t*_182_ = 6.56, *p*_*adj*_ < 0.001, *d* = 0.65; ED-repeat, *t*_182_ = 10.04, *p*_*adj*_ < 0.001, *d* = 1.00; ED-ID, *t*_182_ = 3.59, *p*_*adj*_ < 0.001, *d* = 0.35), whereas the old age group did not showcase a significant difference between any conditions (p≥.057). Additionally, older adults exhibited lower theta power of the early component compared to the young in both the ID (*t*_182_ = 2.48, *p*_*adj*_ = 0.014, *d* = 0.37) and ED condition (*t*_182_ = 3.23, *p*_*adj*_ = 0.002, *d* = 0.48), but not in repeat trials (*t*_182_ = 1.47, *p*_*adj*_ = 0.142, *d* = 0.22) (**Figure 3**B).

**Figure 3:**
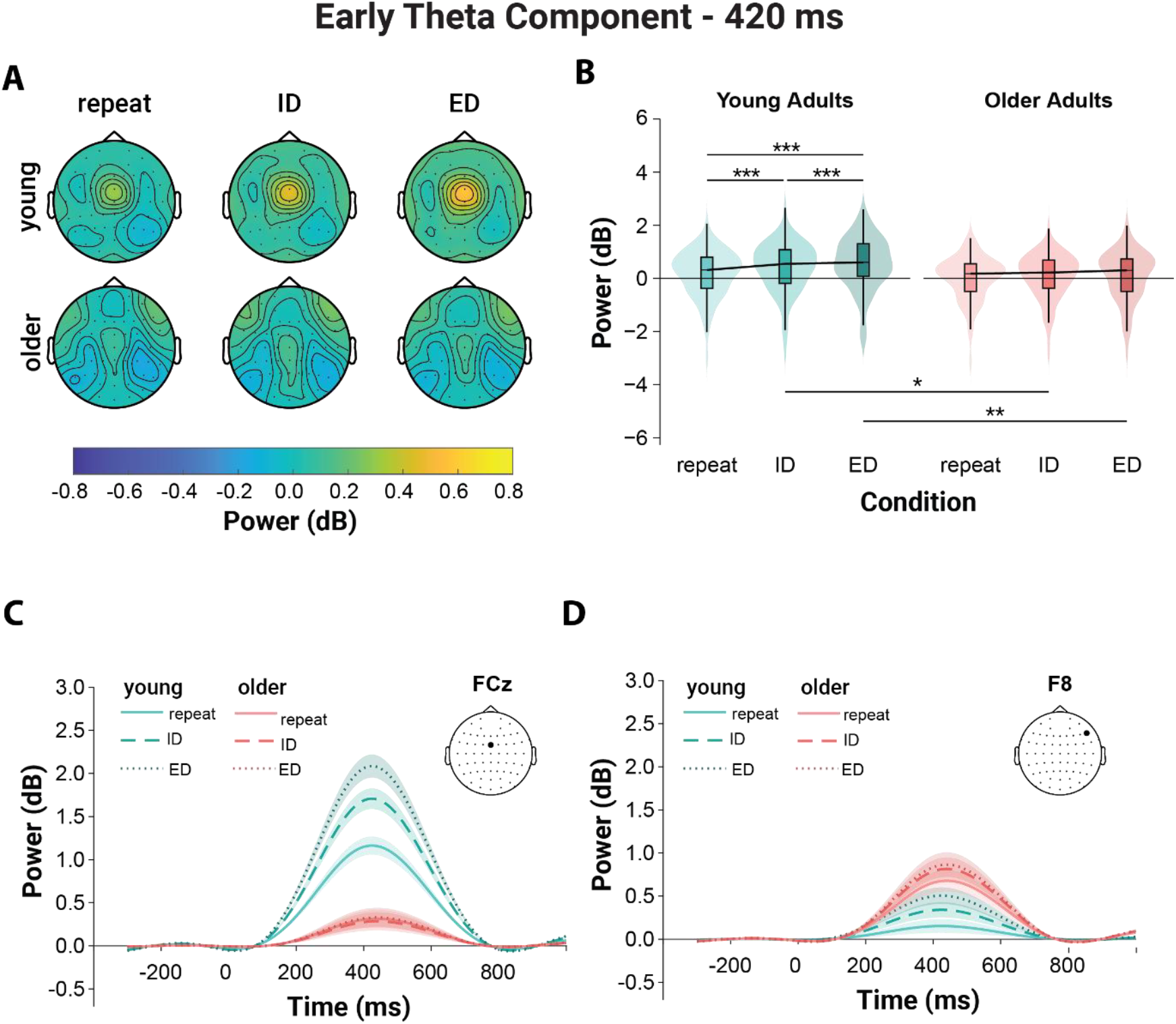
Early Theta Power Component. **A:** Average topographical distribution of the early theta power component at the peak latency of 435 ms. The maximum amplitude of theta at peak latency was at the FCz in the young group and at the F8 in the older age group; **B:** Average component power at peak latency across all electrodes. Young adults display a significant increase in theta power from repeat to ID and ED condition (maximum *t*-value, ED-repeat, *t*_219_ = 11.26, *p*_*adj*_ < 0.001, *d* = 1.32). Older adults display significantly lower theta power then young in both set-shifting conditions (highest *t*-value in ED: *t*_219_ = 3.69, *p*_*adj*_ < 0.003, *d* = 0.50); **C&D:** Theta power timeline of the early theta component at the FCz and F8 channels respectively.

In the young group, the late as the early component peaked over FCz around 680 ms, explaining 27.74% of the variance. In the older age group, the maximum peak of the late theta component was visible over F7 around 690 ms and explained 29.79% of the variance (**Figure 4**). The ANOVA did not reveal a significant interaction between the factors group and condition (*F*_1.60,290.92_ = 2.35, *p* = 0.109, η_p_^2^ = 0.01). A significant effect of condition was found (*F*_1.60,290.92_ = 4.66, *p* = 0.002, η_p_^2^ = 0.02). Post-hoc *t*-tests revealed significant theta power differences between repeat and ED (*t*_182_ = −2.30, *p*_*adj*_ = 0.045, *d* = 0.34), as well as between ID and ED (*t*_182_ = −2.46, *p*_*adj*_ = 0.044, *d* = 0.36), with ED reaching higher theta power values in both comparisons. No significant effect of age group was found (*F*_1,182_ = 0.02, *p* = 0.895, η_p_^2^ < 0.01) (**Figure 4**B).

**Figure 4:**
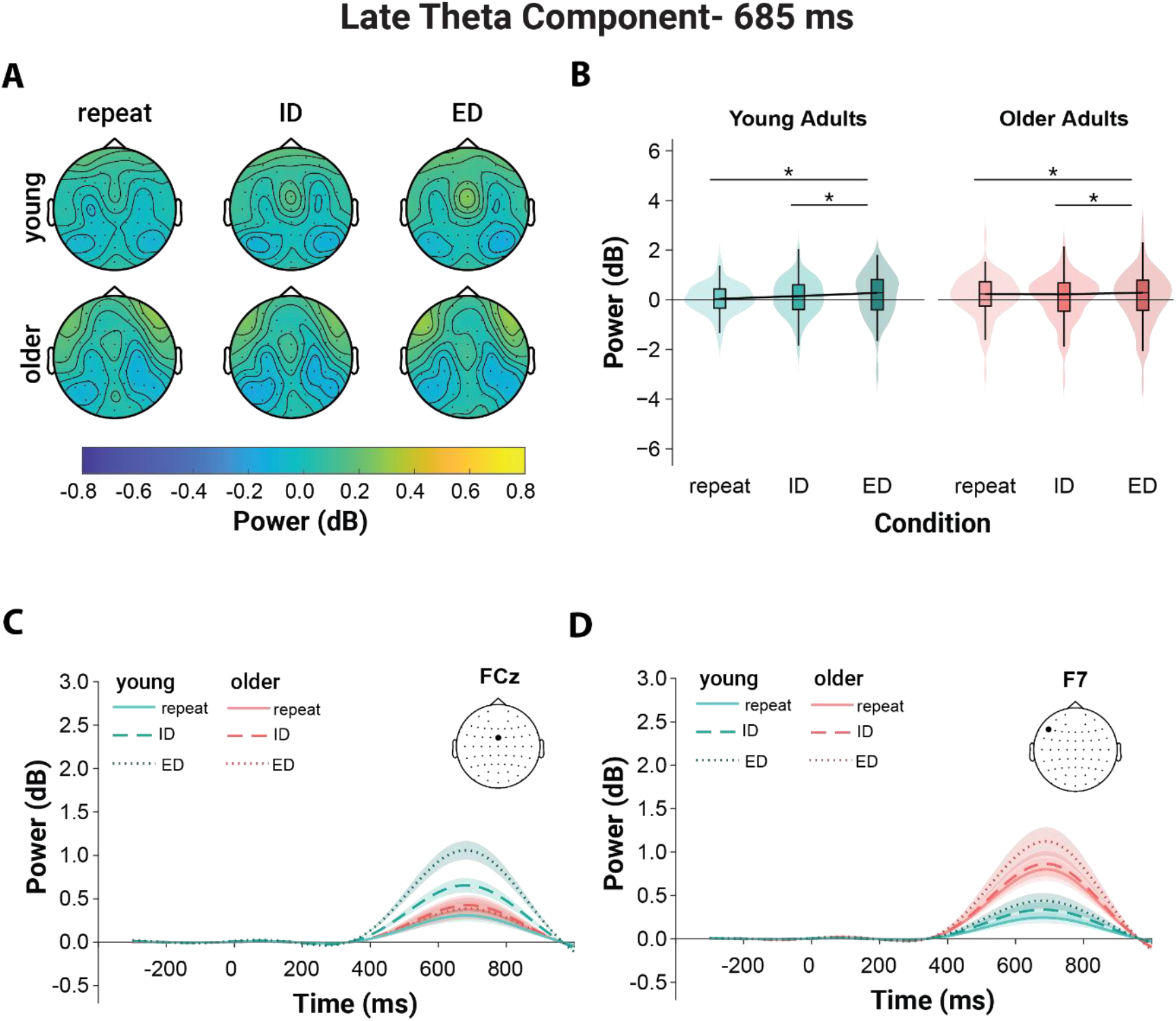
Late Theta Power Component. **A:** Topographical distribution of the late theta power component at the peak latency of 690 ms. The maximum theta power peak was located at FCz in the young age group and at F7 in the older age group; **B:** Average power of the late theta component at peak latency across all electrodes. There was a significant condition effect, in which theta power was higher in the ED condition compared to both the repeat (*t*_219_ = 2.59, *p*_*adj*_ = 0.021, *d* = 0.35) and ID condition (*t*_219_ = 2.91, *p*_*adj*_ = 0.012, *d* = 0.39); **C&D:** Theta power timeline of the late theta component at FCz and F7 respectively.

### 3.3 Theta Coherence

Theta coherence across time was decomposed into factors as described in the previous section. Here, we again identified two components of interest, whose latency peaks reflected the previous components.

The early coherence component reached a peak amplitude at 500 ms in the young age group and explained 12.66% of the variance. The corresponding component in the older age group peaked at 490 ms and explained 11.16% of the variance (**Figure 5**). The ANOVA on the reconstructed components averaged across all channels revealed significant effects of condition (*F*_1,163_ = 6.58, *p* = 0.011, η_p_^2^ = 0.04) and age group (*F*_1,163_ = 7.64, *p* = 0.006, η_p_^2^ = 0.04). Post-hoc tests revealed that theta coherence was higher in the ED condition compared to the ID condition (*t*_163_ = 2.56, *p*_*adj*_ = 0.011, *d* = 0.40). Younger adults displayed overall higher coherence values compared to older adults (*t*_163_ = 2.76, *p*_*adj*_ = 0.006, *d* = 0.43), but no significant interaction between condition and age group was found (*F*_1,163_ = 2.71, *p* = 0.102, η_p_^2^ = 0.02). Importantly, frontocentral electrodes were strongly represented in the electrode pairs belonging in the 95^th^ percentile of the component peak, mirroring the theta-power component peak location (**Figure 5**). This was not the case in the older age group, as here no specific focal location was apparent in the topography.

**Figure 5:**
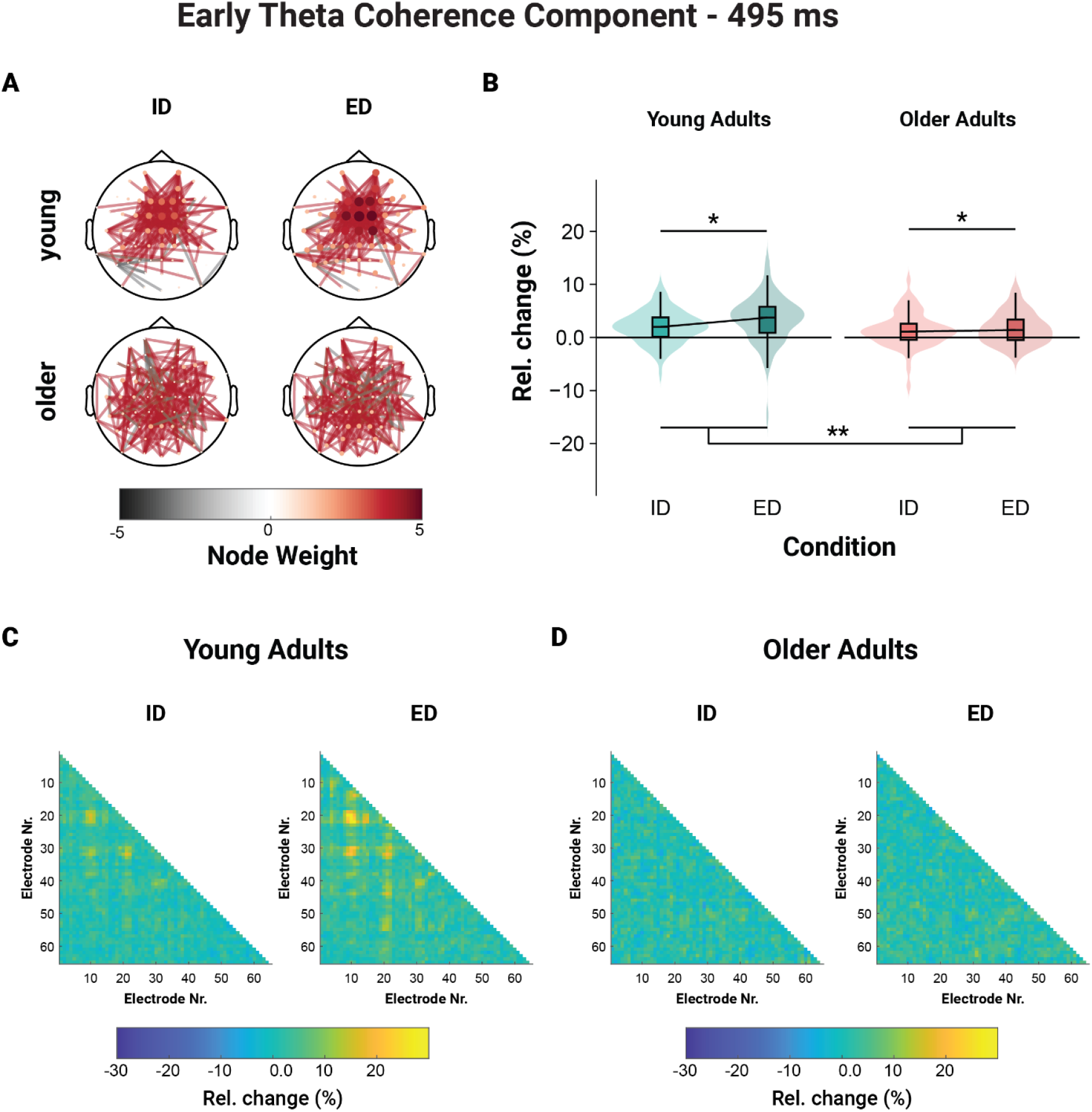
Early Theta Coherence Component. **A:** Connectivity graph of the electrode pairs with absolute score values higher than the 95^th^ percentile; **B:** Average relative change in coherence from repeat at peak latency across all electrode pairs. There was a significant condition effect, in which relative coherence change was higher in the ED condition compared to ID (*t*_163_ = 2.56, *p*_*adj*_ = 0.011, *d* = 0.40). Additionally, younger adults displayed a higher overall relative change compared to older adults (*t*_163_ = 2.76, *p*_*adj*_ = 0.006, *d* = 0.43); **C&D:** Representation of relative change from repeat in the individual conditions in young and old adults respectively.

The late coherence component reached a peak amplitude at 650 ms in the young age group and explained 11.70% of the variance. The equivalent component in the older age group reached a peak at 640 ms and explained a variance of 11.75%. Here, we found no significant effect of age group, condition or their interaction (p ≥ .295).

### 3.4 Global efficiency

We investigated global efficiency in two separate time windows, reflecting the timing of the theta power and coherence peaks described in the previous sections. In the early window from 410 ms to 510 ms, we found a significant interaction between age group and condition (*F*_1.79,289.97_ = 5.46, *p* = 0.003, η_p_^2^ = 0.03) (**Figure 6**A). Young adults exhibited increased global efficiency in both ID (*t*_97_ = 5.96, *p*_*adj*_ < 0.001, *d* = 0.61) and ED compared to repeat trials (*t*_97_ = 6.31, *p*_*adj*_ < 0.001, *d* = 0.64) and during ED compared to ID trials (*t*_97_ = 2.73, *p*_*adj*_ = 0.008, *d* = 0.28). On the other hand, older adults displayed significantly higher global efficiency in ID compared to repeat (*t*_67_ = 2.91, *p*_*adj*_ = 0.010, *d* = 0.36), and ED compared to repeat trials (*t*_67_ = 3.41, *p*_*adj*_ = 0.003, *d* = 0.42) but not in ED compared to ID trials (*t*_67_ = 0.06, *p*_*adj*_ = 0.95, *d* = 0.01). Pair-wise comparisons between age groups revealed a significant difference in global efficiency in the repeat condition between the two age groups (*t*_97, 67_ = 2.20, *p*_*adj*_ = 0.029, *d* = 0.35), with older adults displaying higher values. However, no significant group difference was found in the two set-shifting conditions (*p* ≥ 0.226).

**Figure 6:**
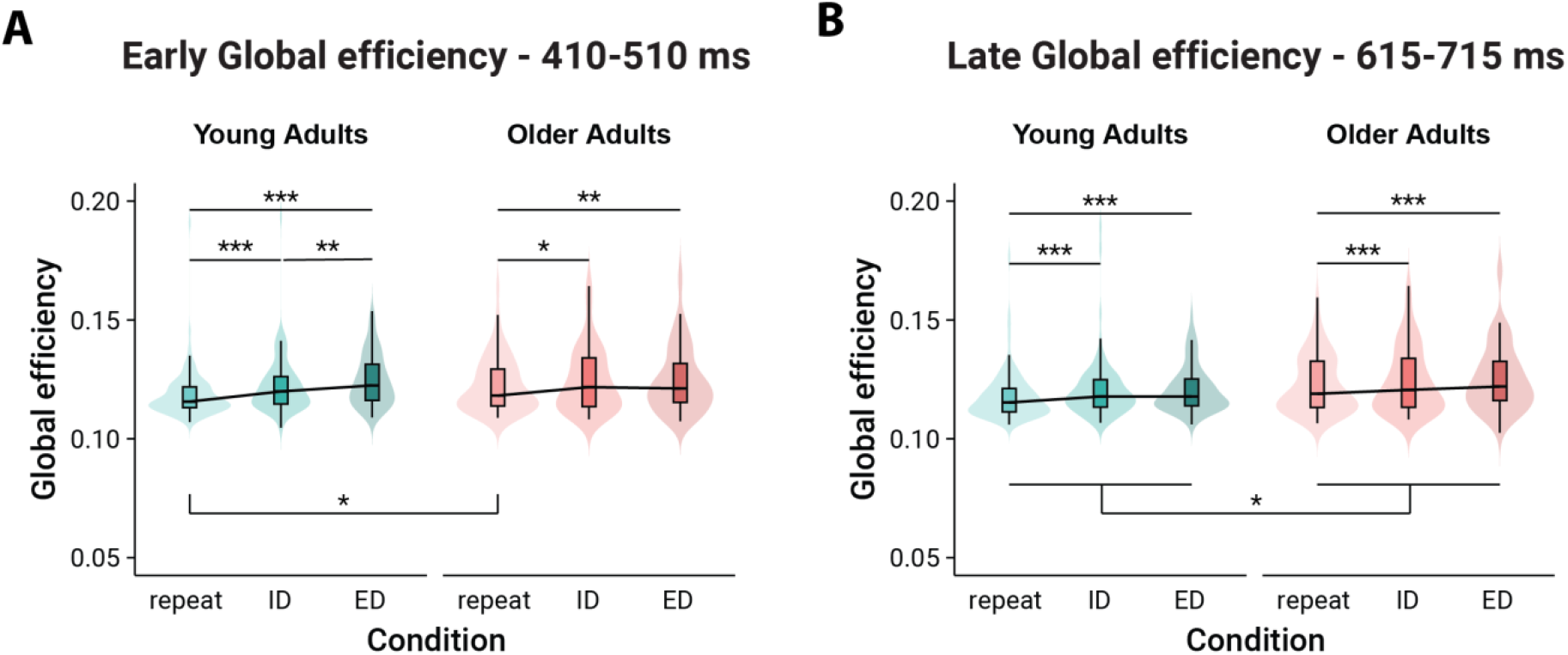
Global efficiency. **A:** Average global efficiency in the early time window between 410 and 510 ms. Young individuals displayed significantly increasing global efficiency from repeat to ID and ED and older adults showed an increase from repeat to ID and from repeat to ED (highest *t*-value in ED-repeat: *t*_97_ = 6.31, *p*_*adj*_ < 0.001, *d* = 0.64). Additionally, younger adults displayed a lower global efficiency in the repeat condition compared to the older participants (*t*_97, 67_ = 2.20, *p*_*adj*_ = 0.029, *d* = 0.35); **B:** Average global efficiency in the late time window between 615 and 715 ms. There was a significant condition effect, in which both set-shifting displayed increased global efficiency compared to repeat (highest *t*-value in ED-repeat: *t*_164_ = 4.44, *p*_*adj*_ < 0.001, *d* = 0.35). Additionally, younger adults displayed a lower global efficiency than older adults (*F*_1.79,290.17_ = 0.29, *p* = 0.727, η_p_^2^ = 0.00).

**Figure 7:**
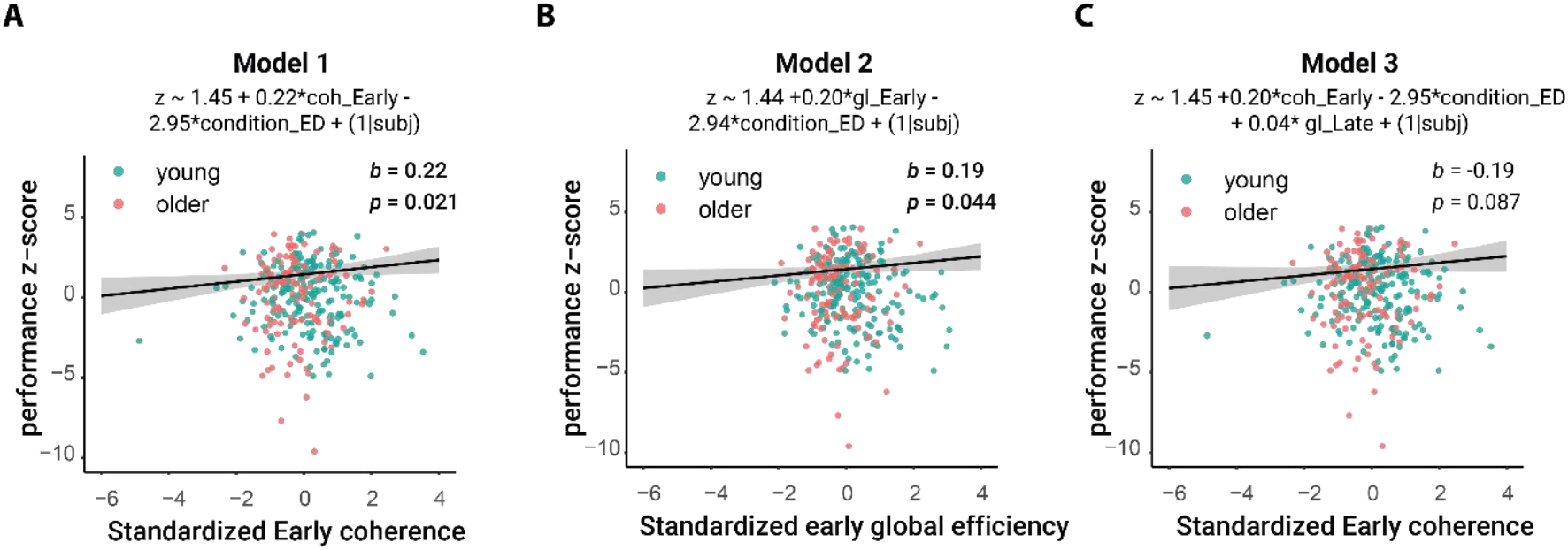
Models to predict performance. **A:** Estimated regression between standardized early coherence and performance z-score obtained from the first model fitting the data (b = 0.22, SE = 0.10, *t*_286.49_ = 2.33, *p* = 0.021); **B:** Estimated regression between standardized early global efficiency and performance z-score obtained from the second model fitting the data (b = 0.19, SE = 0.10, *t*_293.58_ = 2.01, *p* = 0.044); **C:** Estimated regression between standardized early early coherence and performance z-score obtained from the third model fitting the data. Here, the relationship of coherence to z-score is displayed (b = 0.20, SE = 0.12, *t*_296.67_ = 0.38, *p* = 0.087).

In the second time window (615 ms to 715 ms), we found a significant effect of condition (*F*_1.79,290.17_ = 11.07, *p* < 0.001, η_p_^2^ = 0.06), with higher global efficiency in both set-shifting conditions compared to the repeat condition (ID-repeat: *t*_164_ = 4.38, *p*_*adj*_ < 0.001, *d* = 0.34; ED-repeat: *t*_164_ = 4.44, *p*_*adj*_ < 0.001, *d* = 0.35)(**Figure 6**B). We also observed a significant age group effect (*F*_1.79,290.17_ = 6.20, *p* = 0.014, η_p_^2^ = 0.04). Young participants showed lower global efficiency than the older ones. Yet, no significant interaction between condition and age group was found (*F*_1.79,290.17_ = 0.29, *p* = 0.727, η_p_^2^ = 0.00).

### 3.5 Relationship between behaviour and theta measures

The best fitting models describing the relationship between behaviour and theta measures are being described in Table 3.

**Table 3.**
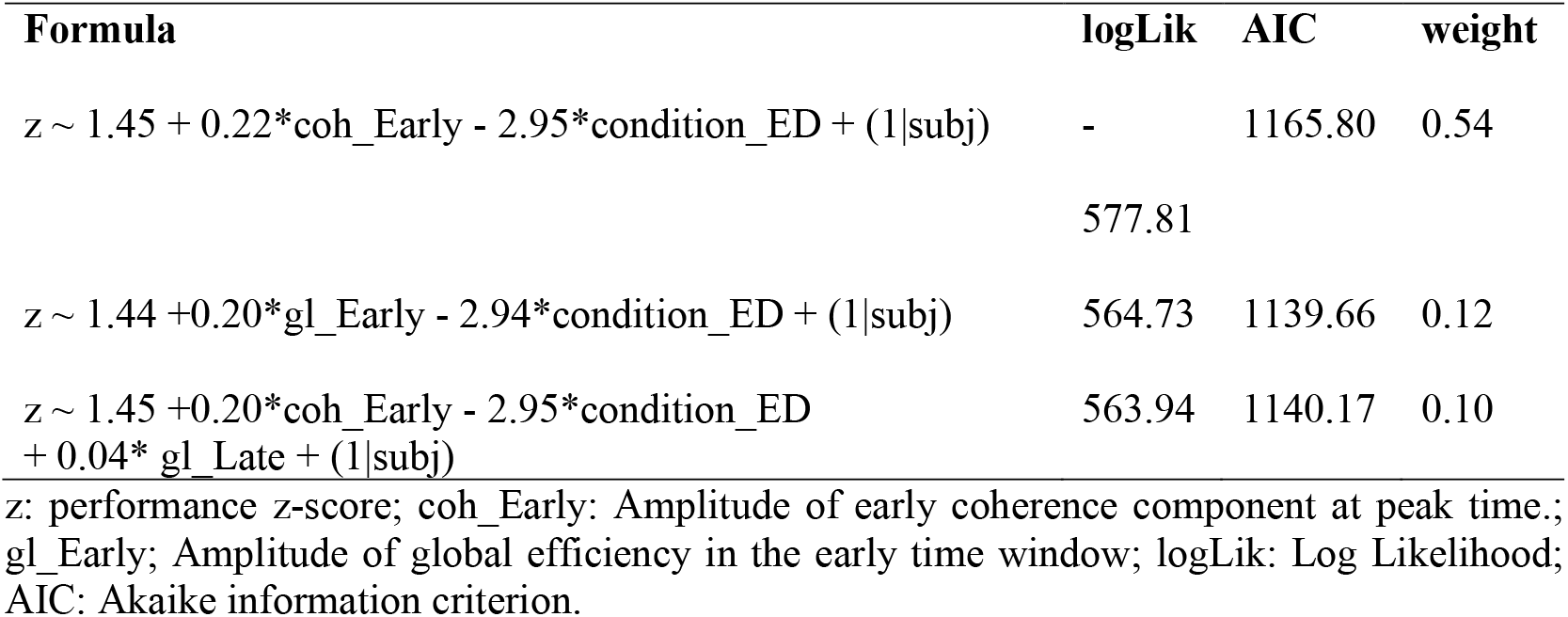
Best fitting linear mixed-effect models.

The best fitting model explained 70.3% of the variance and the early coherence component was a significant predictor of performance (b = 0.22, SE = 0.10, *t*_286.49_ = 2.33, *p* = 0.021), indicating that higher coherence was associated with lower z scores, translating to better performance.

The second-best model explained 70.6% of the variance. Here, it was rather the early global efficiency component that acted as significant predictor of performance with a positive slope (b = 0.19, SE = 0.10, *t*_293.58_ = 2.01, *p* = 0.045), indicating that higher global efficiency values are associated with better performance.

The third best model explained 70.2% of the variance and included both the early coherence component and late global efficiency as predictors with no significance (early coherence: b = 0.20, SE = 0.12, *t*_297.20_ = 1.72, *p* = 0.087; late global efficiency: b = 0.04, SE = 0.12, *t*_296.67_ = 0.38, *p* = 0.708). The direction of the effect here was positive and thus consistent with the slope direction of the previous models.

## 4. Discussion

In this study we aimed to identify whether EEG measures of theta activity and synchrony during set-shifting are modulated differently in older compared to young adults. Our results overall indicated that behaviourally, older adults show a more conservative response pattern with lower error rates and longer reaction times during set-shifting, accompanied by pronounced age differences in the underlying electrophysiological processes.

### 4.1 Speed-accuracy trade-off in set-shifting

As summarized in the introduction, it has thus far been unclear if and how set-shifting performance declines with age. In the present study, using the IDED task, we observed increased RT and IQR costs in older adults, particularly in the ED condition, the most cognitively demanding condition, compatible with the frequently observed age-related slowing of cognitive processes (Salthouse, 1996). From reaction times alone, one might therefore conclude that older adults exhibit behavioural costs in set-shifting mirroring the results of several previous studies (e.g. Cepeda et al., 2001; Kray et al., 2002; Meiran et al., 2001). Conversely, when solely relying on error rates as performance measure, one might conclude that older adults exhibited reduced error switch costs compared to young adults in both set-shifting conditions. However, when we computed z-scores reflecting the composite of all behavioural measures, we found that the global set-shifting ability of older adults was not significantly different from that of younger participants, suggesting an age-related shift in response dynamics. This shift can be summarised as the speed-accuracy trade-off, a phenomenon observed in older age already over four decades ago (e.g. Rabbitt, 1979; Salthouse, 1979). Such a speed-accuracy trade-off aligns with previous research demonstrating similar phenomena in relation to other cognitive tasks, where increased deliberation in older adults may offset declines in processing efficiency (Ratcliff et al., 2003; Starns and Ratcliff, 2010), thus reflecting a compensatory mechanism. As highlighted by Starns and Ratcliff (2010), older adults show a tendency to primarily focus on error minimization, even when speed costs are substantial. This more cautious approach to task-solving is most probably involuntary and may help attenuate adverse effects of a slower processing system (Eckert, 2011; Preprint: Heathcote et al., 2022) or may reflect age-related changes in brain connectivity (Forstmann et al., 2011) (see below).

It cannot be excluded that, in more difficult tasks, older adults may show actually lower performance rather than the speed-accuracy trade-off observed in the IDED task. For example, when participants of our study performed the Flexibility subtask of the TAP, older adults showed both higher error rates and longer reaction times (Table 2). A majority of older adults also failed the ED stage of the ASST task. Those that successfully completed it, required on average more trials than young participants to reach the completion criterion (Table 2).

More generally, our results underscore the importance of considering multiple performance metrics when evaluating age-related cognitive changes, as, for example, relying on reaction time alone may obscure the adaptive strategies employed by older adults to maintain – or, as in our study, even improve – performance accuracy.

### 4.2 Brain connectivity in older age

At the neural level, we observed marked age-related differences in both theta power and connectivity measures, which were differentially modulated during set-shifting in older compared to young adults.

First, we showed that theta modulations do take place during set-shifting in older age, opposing our previous results with a smaller cohort (Darna et al., 2025). This modulation was visible in the ED condition and had its maximum over frontal channels, contrasting that of young adults, which occurred earlier and was most pronounced over frontocentral channels. As the EEG is only a projection of cortical activity, the precise source of this modulation cannot be conclusively determined by our present data, but the overall topography is nevertheless in line with the previously described posterior-anterior shift in aging (Davis et al., 2008; Festini et al., 2018), which refers to the phenomenon that increased frontal activity accompanies a reduction of posterior activity in aging, possibly reflecting functional compensation.

Second, we could show small coherence modulations during set-shifting in older adults. However, the distribution of this modulation was widespread across the scalp, contrasting that of young adults, where increased coherence was most prominent over frontocentral electrodes, in line with previous results (Myers et al., 2021; Sauseng et al., 2006). We attribute this global decrease of coherence modulation to age-related network dedifferentiation (Fjell et al., 2016; Koen et al., 2020) and the reduced specificity of functional connectivity in the aging brain (Geerligs et al., 2014), resulting in decreased within-network connectivity. Increased age-related network dedifferentiation is also supported by our global efficiency analysis, in which values were higher in older adults during set-shifting in both time windows and in all conditions in the late time window. This supports the conclusion that processes associated with focused activity in younger age, shift to a more global response in older adults, where we observe an increase in global activity due to the decreased functionality of the region originally responsible for a specific function, prompting the need for compensatory, more widespread activity. Alternatively, or likely additionally, more global activity in aging may result from reduced inhibition of brain activity not required for the task at hand (Legon et al., 2016; Morcom and Henson, 2018; Schott et al., 2023)-

Hence, our results point to an altered theta distribution and coherence as a neural manifestation of cognitive flexibility in older age. These alterations might explain the speed-accuracy trade-off in older adults and may, at least in part, reflect a compensatory mechanism by which additional cognitive resources are engaged to enable successful set-shifting in older age.

### 4.3 Relationship between EEG measures and behaviour

In a model comparison using the AIC, we found that theta coherence and global efficiency were the EEG measures most robustly associated with performance in our cohort. This is in line with previous studies, in which theta coherence was associated with more favourable indices of cognitive flexibility in older adults (Ferreira et al., 2013). Our results therefore provide further evidence for the possibility that theta coherence and global efficiency may be markers of neural processes underlying set-shifting performance or, more broadly, cognitive flexibility and potentially other executive functions. In line with this interpretation, increased theta-coherence in particular has previously been associated with better performance in tests of executive function (Basharpoor et al., 2021; Dias et al., 2015) and working memory (Smit et al., 2023). Notably, as in the present study, this relationship appears to be independent of age-group (Papenberg et al., 2013). Thus, if these connectivity measures can indeed be regarded as mediators of executive functioning (Li et al., 2020), this opens the possibility of developing targeted tools and interventions aimed specifically at these mechanisms. Such interventions would focus on modulating the neural dynamics reflected in these metrics, for example by increasing theta-band coherence among regions implicated in cognitive control or by enhancing the overall efficiency of large-scale brain networks that support flexible, goal-directed behaviour. Early work in this direction has already yielded promising results, demonstrating enhanced adaptive behaviour following non-invasive brain stimulation (Reinhart, 2017) as well as after neurofeedback training (Enriquez-Geppert et al., 2014).

### 4.4 Conclusion and future perspectives

In summary, our study reveals that set-shifting in older age exhibits a speed-accuracy trade-off, that is, older adults focus on reducing errors at the expense of response speed. This strategy shift may reflect a compensatory mechanism in the presence of age-related functional alterations in the connectivity within and between brain networks. This observation reveals the ability of the aging brain to employ alternative neural resources to compensate for the decline of task-specific neural resources in the form of cognitive reserve, as it is mediated by theta coherence and global efficiency. Prospectively, targeted training protocols and interventions such as non-invasive brain stimulation may help to counteract cognitive decline with or without the presence of a neurodegenerative disease.

## Acknowledgments

The authors are grateful to Matthias Deliano and Renate Blobel-Lüer for their expert technical assistance in the EEG and the students that aided in participant recruitment and data collection (Jascha Jauch, Karla Reif, Leonie Schenk, Beatrice Christin Guhl, Leonie Kaulke and Chi-Hsuan Lu). Finally, we would like to express our sincere gratitude to our study participants for their consent to take part in this research study.

## Author Contribution

MD: conceptualization, data curation, methodology, software, formal analysis, investigation, writing – original draft, writing – review and editing, visualization

AR: conceptualization, supervision, methodology, writing – original draft, writing – review and editing

JMH: methodology, writing – review and editing

CIS: conceptualization, supervision, funding acquisition, writing – original draft, writing – review and editing

BHS: conceptualization, supervision, funding acquisition, writing – original draft, writing – review and editing

## Data Availability Statement

Experimental Code, MATLAB Scripts, RStudio Scripts can be provided upon request.

