## Supplementary Material for "Altered theta distribution and coherence during set-shifting in older age"

### 1 Supplementary Methods

#### 1.1 Neuropsychological Tests

A subset of the recruited cohort was further evaluated with a number of common psychometric tests. In short, we conducted the Verbal Learning and Memory test (VLMT; Helmstaedter et al., 2001)), the alertness subtest from the Test Battery for Attention (TAP; Zimmermann & Fimm, 1992), a flanker task (Eriksen & Eriksen, 1974) and an n-back task (Gevins & Cutillo, 1993). The psychological constructs derived from each test are being described in Table S1, together with the results from each age group. Outliers with values higher than the 3^rd^ quartile + 3*IQR or lower than the 1^st^ quartile – 3*IQR were excluded from the individual analyses (see Table S2 for outlier borders).

**Table S1. Neuropsychological testing battery**

|  |  | **Young**  **M + SD [N]** | **Older**  **M + SD [N]** | **Statistics** |
| --- | --- | --- | --- | --- |
| **Verbal Learning and Memory Test (VLMT)** |  |  |  |  |
| Repetitions of list A (sum score) | Learning ability | 66.08 ± 6.39 [50] | 56.45 ± 10.31 [62] | ***W* = 654.5,**  ***p <* .001** |
| Distractor list B | Pro-active inhibition | 10.02 ± 2.63 [50] | 6.91 ± 2.17 [65] | ***W* = 573,**  ***p <* .001** |
| Recall of list A | Retro-active inhibition | 14.04 ± 1.43 [50] | 11.82 ± 2.60 [62] | ***W* = 676.5,**  ***p <* .001** |
| 30-min delayed recall of list A | Episodic memory | 14.34 ± 0.94 [50] | 11.78 ± 2.64 [65] | ***W* = 612,**  ***p <* .001** |
| one-day delayed recall of list A | Episodic memory | 13.82 ± 1.43 [39] | 9.41 ± 2.99 [49] | ***W* = 181.5,**  ***p <* .001** |
| **Altertness subtest from the TAP** |  |  |  |  |
| RT in trials with cue tone (ms) | Tonic alertness | 239.29 ± 32.08 [50] | 306.39 ± 58.74 [62] | ***W* = 2633,**  ***p <* .001** |
| RT in trials without cue tone (ms) | Phasic altertness | 264.09 ± 30.41 [50] | 342.09 ± 55.88 [62] | ***W* = 2796.5,**  ***p <* .001** |
| **Flanker task** |  |  |  |  |
| RT difference (incongruent - congruent | Interference processing | 103.63 ± 53.96 [52] | 206.65 ± 134.71 [57] | ***W* = 2336,**  ***p <* .001** |
| **N-Back task** |  |  |  |  |
| 1-back corrected hit rate (%) | Working memory | 97.29 ± 4.34 [53] | 91.10 ± 11.20 [59] | ***W* = 1047,**  ***p <* .001** |
| 1-back RT (ms) | Working memory | 440.95 ± 58.80 [53] | 502.31 ± 78.77 [59] | ***W* = 2313,**  ***p <* .001** |
| 2-back corrected hit rate (%) | Working memory | 68.99 ± 23.73 [53] | 19.17 ± 33.79 [59] | ***W* = 318.5,**  ***p <* .001** |
| 2-back RT (ms) | Working memory | 590.43 ± 82.39 [53] | 697.63 ± 107.11 [59] | ***W* = 2449,**  ***p <* .001** |
| 3-back corrected hit rate (%) | Working memory | 25.24 ± 34.69 [53] | -9.53 ± 32.17 [59] | ***W* = 706.5,**  ***p <* .001** |
| 3-back RT (ms) | Working memory | 652.85 ± 99.31 [53] | 699.48 ± 142.02 [59] | *W* = 1883,  *p =* .063 |

M: mean; SD: standard deviation; N: number of outliers; VLMT: Verbal Learning and Memory Test (Helmstaedter et al., 2001); TAP: Test Battery for Attention (Zimmermann & Fimm, 1992).

#### 1.3 Outlier detection

**Table S2. Variables and borders for removal of extreme outliers in young and older participants**

|  | **Young**  **Criterion [N]** | **Older**  **Criterion [N]** |
| --- | --- | --- |
| **IDED** |  |  |
| Reaction Time (ms) | > 640.59 [2] |  |
| IQR | > 291.71 [2] | - |
| Error rate (%) | - | > 9.33 [1] |
| **Flexibility subtest from the TAP** |  |  |
| Error rate (%) | > 24.92 [1] | - |
| RT (ms) | - | - |
| **Verbal Learning and Memory Test (VLMT)** |  |  |
| Repetitions of list A (sum score) | - | - |
| Distractor list B | - | - |
| Recall of list A | < 7 [1] | - |
| 30-min delayed recall of list A | < 11 [1] | - |
| one-day delayed recall of list A | - | - |
| **Altertness subtest from the TAP** |  |  |
| RT in trials with cue tone (ms) |  |  |
| accuracy in trials with cue tone (%) | < 90.00 [2] | < 90.00 [3] |
| RT in trials without cue tone (ms) |  |  |
| accuracy in trials without cue tone (%) | < 97.00 [2] | < 97.00 [2] |
| **Flanker task** |  |  |
| accuracy in congruent trials (%) | < 91.25 [2] | < 90.00 [3] |
| RT in congruent trials (ms) | - | > 1069.83 [1] |
| accuracy in incongruent trials (%) | < 28.44 [1] | < 30.00 [2] |
| RT in incongruent trials (ms) | - | > 1518.64 [2] |
| **N-Back task** |  |  |
| 1-back corrected hit rate (%) | < 75.00 [1] | < 25.00 [6] |
| 1-back RT (ms) | - | > 826.16 [5] |
| 2-back corrected hit rate (%) | - | - |
| 2-back RT (ms) | - | - |
| 3-back corrected hit rate (%) | - | < - 162.50[1] |
| 3-back RT (ms) | - | > 1343.39 [1] |

Classification as extreme outliers based on the interquartile range (x > 3rd quantile + 3*interquartile range or x < 1st quantile - 3*interquartile range); N: number of outliers; TAP: Test Battery for Attention (Zimmermann & Fimm, 1992). VLMT: Verbal Learning and Memory Test (Helmstaedter et al., 2001).

#### 1.4 Attentional Set Shifting task (ASST)

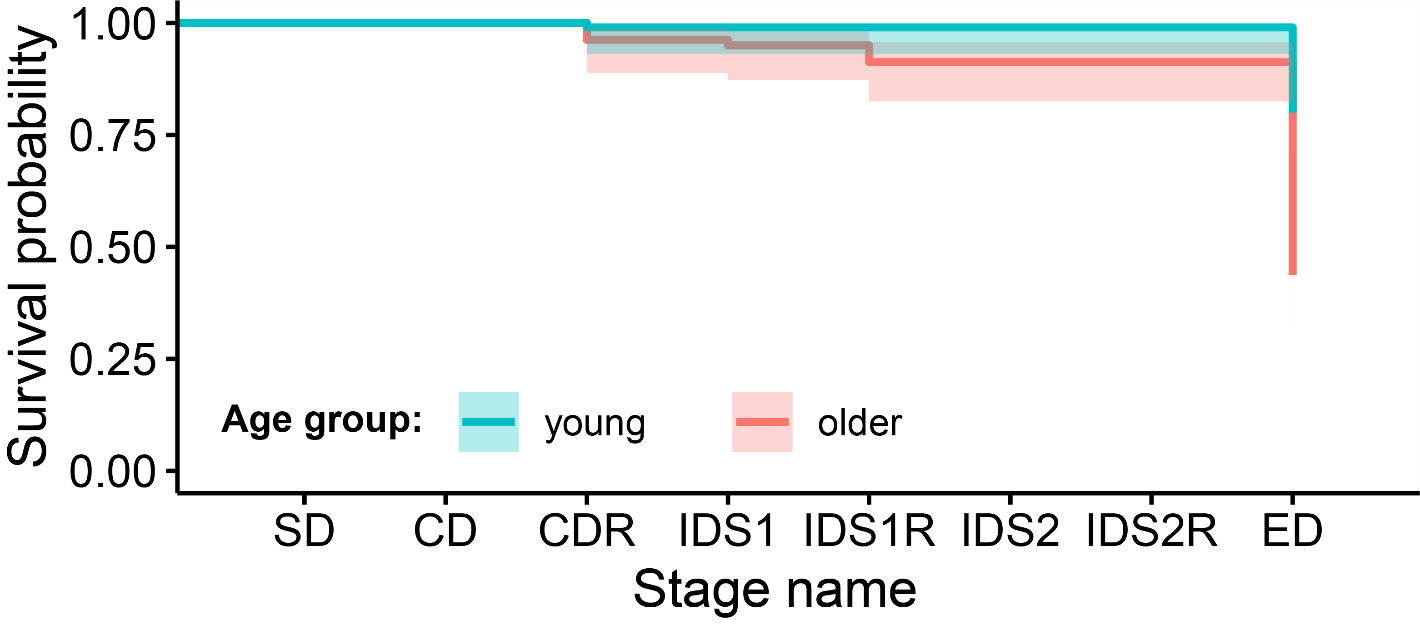

**Supplementary Figure 1:** Probability to complete the Attentional Set Shifting Task (ASST) for the two age groups, plotted as survival curve. There is a significant difference in task completion rates between the two age groups (X^2^(1, N = 180) = 26.40, p < .001). The majority of older individuals fail to complete the ASST at the ED stage. The shaded areas represent Standard Errors of the survival curves. SD: Simple Discrimination; CD: Compound Discrimination; CDR: Compound Discrimination Reversal; IDS1: Intra-dimensional shift 1; IDS1R: Intra-dimensional shift 1 reversal; IDS2: Intra-dimensional shift 2; IDS2R: Intra-dimensional shift 2 reversal; ED: Extra-dimensional shift.

#### 1.5 Artifact detection and correction

We used the EEGLAB plug-in microDetect (Craddock et al., 2016) to detect and correct microsaccadic artifacts for the purposes of investigating gamma-band range (Keren et al., 2010). Microsaccades were detected in epochs of -400 to 700 ms around the stimulus, in trials without blinks using an adaptive threshold and filtered data using a 6^th^ order bandpass Butterworth filter from 30 Hz to 100 Hz. Microsaccade epochs were then built by only evaluating the peri-stimulus period from 100 to 700 ms as suggested in Keren et al. (2010), time locked at -30 ms to 50 ms around each microsaccade. Both the mean centered trial epochs and microsaccade epochs were included in an Independent Component Analysis (ICA) to detect components of blinks, saccades and microsaccades (FASTICA; <http://research.ics.aalto.fi/ica/fastica/>). We then calculated the Pearson correlation coefficient between the individual components and the data obtained from the EOG channels and removed the components with r > 0.33 with any of the 3 EOG channels. We also removed components that reached r > 0.33 with the mastoid channels TP9 and TP10, as these represent muscle artifacts.

#### 1.6 Channel Interpolation

**Table S3. Interpolated channels followed by the respective number of subjects for which the interpolation was performed separated by age group.**

| **Channel** | **No. of young subjects** | **No. of older subjects** |
| --- | --- | --- |
| FC5 | - | 3 |
| T8 | - | 4 |
| TP8 | 1 | - |
| P5 | 1 | - |
| POz | 1 | - |

### 2 Supplementary Results

#### 2.1 Raw behaviour results

**Table S4. IDED Behavioural results in young and older participants**

|  | **Young**  **M + SD** | **Older**  **M + SD** |
| --- | --- | --- |
| **Repeat** |  |  |
| Median RT (ms) | 433.05 ±37.64 | 651.86 ±105.34 |
| IQR (ms) | 122.05±25.41 | 200.59±51.05 |
| Error rate (%) | 2.77 ± 2.35 | 1.27 ±1.30 |
| **ID** |  |  |
| Median RT (ms) | 494.30 ±42.99 | 728.29 ±109.66 |
| IQR (ms) | 149.92±31.55 | 232.53±59.20 |
| Error rate (%) | 6.31 ±5.20 | 2.02 ±1.88 |
| **ED** |  |  |
| Median RT (ms) | 553.90 ± 61.72 | 841.23 ±147.63 |
| IQR (ms) | 196.11±47.32 | 303.36±80.51 |
| Error rate (%) | 9.76±5.80 | 4.07±3.42 |
